# The crabeater seal reference genome reveals hallmarks of persistently large effective population size and sustained population expansion in the World’s most abundant pinniped

**DOI:** 10.64898/2026.06.16.732718

**Authors:** Beril Yıldız, Thomas Gelatt, Luis A. Hückstädt, Daniel P. Costa, Michael Tift, Jay Rotella, Elizabeth Flesch, Kaitlin Macdonald, Nancy Chen, Robert Garrott, Mike E. Goebel, Jaume Forcada, Thorsten Wachtmeister, David L. J. Vendrami, Joseph I. Hoffman

**Affiliations:** Department of Evolutionary Population Genetics, Faculty of Biology, Bielefeld University, 33501 Bielefeld, Germany; Center for Biotechnology (CeBiTec), Faculty of Biology, Bielefeld University, 33615 Bielefeld, Germany; Alaska Fisheries Science Center, National Oceanic and Atmospheric Administration, Seattle, WA, USA; Institute of Marine Science, University of California Santa Cruz, Santa Cruz, California, USA; Centre for Ecology and Conservation, College of Life and Environmental Science, University of Exeter, Penryn, Cornwall, UK; Sharjah Marine Science Research Centre, University of Khorfakkan, UAE; Department of Biology and Marine Biology, University of North Carolina Wilmington, Wilmington, North Carolina, USA; Department of Ecology, Montana State University, P.O. Box 173460, Bozeman, MT 59717, USA; Department of Ecology and Evolutionary Biology, University of California, Los Angeles, California, USA; Antarctic Ecosystem Research Division, Southwest Fisheries Science Center, NOAA Fisheries, La Jolla, California, USA; Institute of Marine Sciences, University of California Santa Cruz, Santa Cruz, California, USA; British Antarctic Survery, High Cross, Madingley Road, Cambridge CB3 OET, UK; Genomics & Transcriptomics Laboratory, Biologisch-Medizinisches Forschungszentrum, and West German Genome Center, Heinrich-Heine-Universität Düsseldorf, Universitätsstr. 1, 40225, Düsseldorf, Germany; Joint Institute for Individualisation in a Changing Environment (JICE), Bielefeld University and University of Münster, Bielefeld, Germany

**Keywords:** genome assembly, crabeater seal, pinniped, effective population size, genomic diversity, inbreeding, population expansion

## Abstract

Population genetic theory predicts that a species’ demographic history shapes patterns of genome-wide variation. However, conservation genomic studies have disproportionately focused on small or declining species, where low genetic diversity and inbreeding are major concerns, while highly abundant species have attracted comparatively less attention. Here, we investigate the crabeater seal (*Lobodon carcinophaga*) which, despite being one of the most numerous large mammals on Earth, remains largely uncharacterised in terms of its genomic diversity and demographic history. We assembled a high-quality crabeater seal reference genome from a combination of Illumina and PacBio HiFi reads, generating a 2.44 Gb assembly spanning 138 scaffolds with high completeness. To evaluate genomic diversity in a comparative context, we whole-genome resequenced 20 crabeater seals alongside 20 individuals each of three Antarctic phocids spanning a population size gradient: the Weddell seal (*Leptopnychotes weddellii*), leopard seal (*Hydrurga leptonyx*) and southern elephant seal (*Mirounga leonina*). Crabeater seals carried 61.5 million SNPs compared to 12–16 million in the other species and exhibited markedly higher nucleotide diversity and negligible genomic inbreeding. We observed an excess of rare alleles, with nearly half of all variants segregating at frequencies below 5%. Demographic reconstruction revealed persistently large effective population sizes over the past million years and sustained population expansion, paralleling inferred increases in Antarctic krill associated with sea-ice expansion during the late Pleistocene. This study provides a new genomic resource and sheds new light on the evolutionary dynamics of the world’s most abundant pinniped.

## Introduction

Population size changes through time, whether expansion during periods of ecological opportunity or decline under environmental constraint, fundamentally shape patterns of genetic variation within species. Such changes alter the balance among mutation, genetic drift and natural selection, leaving distinct and enduring signatures across the genome (Tajima, 1989). For example, demographic expansions typically generate an excess of rare variants as newly arising mutations accumulate faster than they are removed by drift, whereas bottlenecks reduce genetic diversity and shift the site frequency spectrum (SFS) towards intermediate frequency variants (Nei et al., 1975). Consequently, demographic changes structure standing genetic variation in predictable ways, influencing genome-wide diversity and, in turn, adaptive potential (Frankham et al., 1999). Understanding how demographic histories translate into genome-wide patterns of variation is therefore central to evolutionary and conservation genomics.

A key parameter linking demography to genomic diversity is the effective population size (*N*_e_) (Wright, 1931; Frankham, 1995; Charlesworth, 2009). Because nucleotide diversity (π), defined as the average number of pairwise sequence differences among individuals, is expected to scale with the product of long-term *N*_e_ and the mutation rate, μ, such that π = 4*N*_e_μ in diploid organisms (Nei et al., 1975), demographic changes directly influence levels of genetic diversity through their effects on *N*_e_. Consequently, species that have experienced prolonged small *N*_e_ or bottlenecks tend to exhibit reduced π, whereas those with historically large and stable *N*_e_ maintain higher levels of genetic variation.

Small *N*_e_ is also reflected in patterns of genomic inbreeding. As population size declines, the probability that individuals inherit identical by descent (IBD) haplotypes from both parents increases, leading to the accumulation of runs of homozygosity (ROH) across the genome (Ceballos et al., 2018). Elevated homozygosity unmasks the effects of deleterious alleles, causing inbreeding depression, while also exposing them to selection. The fraction of the genome in ROH (*F*_ROH_) quantifies autozygosity and captures the cumulative effects of both recent and historical population size changes (McQuillan et al., 2008). Accordingly, ROH-based metrics are widely used to quantify autozygosity and to evaluate the genomic consequences of past demographic changes in wild populations (Shafer & Kardos, 2025).

Because species with small or declining *N*_e_ are most susceptible to the loss of genetic diversity and inbreeding, empirical studies of inbreeding have often focused on small or declining populations. By contrast, species with historically large and stable populations are expected to maintain higher π and to exhibit reduced inbreeding (Frankham, 1996). However, the relationship between *N*_e_ and genomic diversity is not always straightforward, as genetic diversity varies far less across species than predicted from differences in population size (“Lewontin’s paradox”, (Lewontin, 1974)). This arises because linked selection, variation in mutation rate and demographic changes can decouple population size from empirical patterns of genetic variation (Romiguier et al., 2014; Lynch et al., 2016; Charlesworth & Jensen, 2022). Consequently, the underrepresentation of species with large and stable populations in studies of wild, non-model organisms leaves a gap in our understanding of how demographic history shapes genomic variation across the full spectrum of *N*_e_.

Pinnipeds provide an ideal system to study the relationship between *N*_e_ and genomic diversity (Stoffel et al., 2018; Peart et al., 2020). They vary enormously in population size, ranging from a few hundred Mediterranean monk seals (*Monachus monachus*) to several million crabeater seals (*Lobodon carcinophaga*) and encompass species with markedly different demographic histories. Some were driven to the brink of extinction by commercial sealing, such as the northern elephant seal (*Mirounga angustirostris*), whereas others inhabiting remote regions such as Antarctica experienced little to no human impacts (Mittermeier & Wilson, 2014). To investigate how these factors shape genomic variation across taxa, high-quality genome assemblies are needed to identify ROHs and accurately reconstruct historical changes in *N*_e_ (Brüniche-Olsen et al., 2018; Li & Durbin, 2011). However, despite recent advances and the growing availability of chromosomal reference genomes (Mohr et al., 2022; Hench et al., 2024; Nebenführ et al., 2025), pinniped species inhabiting remote, ice-covered environments remain underrepresented due to logistic challenges such as harsh weather, short breeding seasons and limited access for sampling.

The crabeater seal (*Lobodon carcinophaga*) is the world’s most numerous pinniped and one of the most abundant large mammals globally, with population size estimates varying from fewer than one million to 75 million individuals (Bengtson, 2009; Southwell et al., 2008), although the true population size is probably in the order of four million (Hückstädt, 2025). Occupying pack ice habitats across the Southern Ocean, it was largely protected from the commercial sealing of the 18^th^ and 19^th^ centuries. As a major consumer of Antarctic krill (*Euphausia superba*), it plays a key role in the krill-based food web of the Southern Ocean (Southwell et al., 2008). Despite its ecological importance, however, its genomic diversity and demographic history remain largely unexplored. Although this species has been included in a handful of phylogenetic and genetic studies based on mitochondrial DNA and small numbers of nuclear markers (Curtis et al., 2009, 2011; Davis et al., 2008), little is known about patterns of genome-wide diversity. Consequently, this species provides an opportunity to test whether the signatures of elevated π and reduced inbreeding expected under large *N*_e_ are confirmed at the genomic scale.

Here, we generated a high-quality reference genome for the crabeater seal and resequenced the genomes of 20 individuals. For comparison, we also sequenced 20 genomes each from three other Antarctic phocids: the leopard seal (*Hydrurga leptonyx*), southern elephant seal (*Mirounga leonina*) and Weddell seal (*Leptonychotes weddellii*). These species span a gradient of global population sizes, from ∼220,000–440,000 leopard seals (Rogers, 2009) through ∼750,000 southern elephant seals (Hindell et al., 2016) to ∼202,000 Weddell seals (LaRue et al., 2021). Only the southern elephant seal experienced significant hunting during the 19^th^ century hunting, yet it did not collapse and has since recovered (Kim et al., 2020).

Using whole-genome resequencing data from all four species, we quantified π and *F*_ROH_ and reconstructed historical changes in *N*_e._ We hypothesized that the crabeater seal would exhibit (i) higher genome-wide diversity; (ii) minimal inbreeding; and (iii) historically large and stable *N*_e_ relative to the other Antarctic phocids.

## Materials and methods

### Study sites and sample collection

Sample collection and sequencing procedures for each species are described below.

### Crabeater seal

Whole blood samples were collected from eight females, six males and one individual of unknown sex, along with a skin sample from a male. Blood samples were obtained from the extradural intervertebral vein in the lower lumbar region using an 18-gauge syringe during an icebreaker cruise in 1994 around the Amundsen and Bellingshausen Seas. The skin sample was collected in 1987 in Barne Glacier, McMurdo Sound, Antarctica (Gelatt & Siniff, 1994). These samples were archived in liquid nitrogen-cooled cryovats that maintain vapor-phase nitrogen at -170°C in the University of Alaska Museum (Arctos). Data associated with these samples are publicly available online via the Arctos database (https://arctos.database.museum/). Sampling was conducted in accordance with the Antarctic Conservation Act of 1978 (FR Doc No: 94-662) and under the National Marine Fisheries Service (NMFS) permit 976. An additional four whole blood samples (three females and one male) were collected in 2022 and 2023 from the Gerlache Strait in the western Antarctic Peninsula and stored at -80°C. This sampling was conducted in accordance with the UC Santa Cruz Institutional Animal Care and Use Committee (Costd1808) under NMFS permit 25770. Sampling was granted to LAH, DPC and MST by National Science Foundation (NSF) grant (OPP 2042032).

### Leopard seal

Skin tissue samples were collected from eight males, seven females and six individuals of unknown sex at Bird Island, South Georgia, between 2003 and 2016. Samples were collected from the webbing of the hindflipper using piglet ear notching pliers, then stored in dimethyl sulfoxide (DMSO) saturated with salt at −20 °C. Sampling was conducted in accordance with the Department for Environment, Food and Rural Affairs (DEFRA) (TARP/2016/054, AHZ/2024A/2005/1) and the Office of the Commissioner of South Georgia and the South Sandwich Islands (SGSSI) (SCI/2014/032).

### Southern elephant seal

Skin tissue samples were collected from one adult male and 19 pups from a breeding colony at Half Moon Beach, Cape Shirreff in the South Shetland Islands between 2008 and 2016 (see Nichols et al., 2022 for details). Adults were sampled from the flanks using a 2 mm sterile disposable Miltex biopsy punch (Thermo Fisher Scientific). Pup skin samples were collected from the rear flipper using a tag hole punch or a 2 mm sterile disposable Miltex biopsy punch. The samples were immediately transferred to 95% ethanol and stored at −20 °C. Sampling was conducted in accordance with the MMPA of 1972 (16472-01 and 774-1847-04) under NMFS permits 2012-005 and 2008-008.

### Weddell seal

Skin tissue samples were collected in 2006 from nine male and 11 female pups born that year in Erebus Bay, Ross Island, Antarctica, as part of a long-term demographic monitoring program established in 1969 (Powell et al., 2023; Proffitt et al., 2007; Rotella et al., 2009). The work that provided and curated the samples was supported by a series of grants from the NSF, Office of Polar Programs (Grant Nos. 1640481, 2147553 and 2147554) and prior NSF Grants to R.A. Garrott, J. J. Rotella, D. B. Siniff, and J. Ward Testa. Sampling was conducted in accordance with the Marine Mammal Protection Act of 1972 (MMPA; 16 U.S.C. 1361 et seq.) under NMFS permit 26375.

### Crabeater seal genome assembly

A blood sample from a female juvenile crabeater seal captured in 2023 in the Le Maire Strait, western Antarctica, was used for the genome assembly. High-molecular-weight DNA was extracted using a QIAGEN Genomic-tip 20G kit. Illumina libraries were generated using the Illumina DNA PCR-Free Prep, Tagmentation (Illumina Inc.) following the manufacturer’s instructions. PacBio libraries were constructed following the PacBio HiFi (CCS) standard protocol and sequenced on a PacBio Revio platform by BMKgene. HiFi sequencing generated a total of 106.6 Gb of HiFi data with an average sequencing depth of 51x and an average read length of 12.3 Kb. The raw data were processed for adapter and low-quality data removal using smrtlink (v9.0) CCC program with parameters “–minLength 50 –maxLength 50000 –minPasses 1 – minPredictedAccuracy 0.99 -j 6”. The filtered data contained 8,668,775 reads.

The genome assembly was performed using a hybrid approach in Hifiasm (v.024; Cheng et al., 2021) combining Illumina PE150 reads (BGISEQ-500) and PacBio HiFi reads (Pacbio Revio). Scaffolds shorter than 1,000 bp were excluded. The quality of the assembly was assessed by evaluating conserved gene content with CEGMA (v2.5) using the default parameters, and BUSCO (v5.8.2; options: -m genome -c 24 -e 1e-3 --augustus) using the carnivora dataset from OrthoDB v10 as reference. In addition, the Illumina and PacBio reads were aligned to the assembly to determine the mapping rate, genome coverage and sequencing depth. Illumina reads were aligned with BWA (v0.7.10) using default settings and PacBio reads with minimap2 (v2.14-r883; options: -I 20G --MD - ax asm20 t). The assembly statistics were visualised as a snail plot using blobtoolkit (Challis et al., 2020).

### Whole genome resequencing

We generated whole-genome resequencing data for 20 individuals from each species in this study, targeting a sequencing depth of 25x per individual. Whole genomic DNA was extracted using a standardised phenol-chloroform protocol (Sambrook 2001). The DNA extracts were first assessed photometrically using a NanoDrop One spectrophotometer (Thermo Fisher Scientific Inc.) to determine their purity. DNA integrity was then evaluated using an Agilent 5300 Fragment Analyzer System with the DNF-464 HS Large Fragment 50kb Kit (Agilent Technologies, Inc.) and the final DNA concentration was quantified fluorometrically using a Qubit dsDNA-BR assay (Thermo Fisher Scientific Inc.). Crabeater seal whole-genome sequencing libraries were prepared according to the PCR-free BGI-reseq protocol and 150bp PE sequenced on a BGISEQ-500 by BMKgene. All other samples were prepared using the Illumina DNA PCR-Free Prep Tagmentation kit (Illumina Inc.) according to the manufacturer’s protocol, with >300 ng input DNA per sample. Libraries were normalized to 2 nM, pooled and 151bp PE sequenced on a NovaSeq 6000 system (Illumina Inc.) at the West German Genome Center.

### Bioinformatics and variant calling

Read quality was assessed using FastQC v0.11.9 (Andrews, 2010), checked for potential interspecific contamination using fastq_screen v0.15.1 (Wingett & Andrews, 2018) and visualised using multiQC v1.12 (Ewels et al., 2016). Reads with phred quality scores below 20 (-q 20) were removed using fastp v0.23.4. The remaining reads were then mapped to the respective reference genome (LepWed1.0 for Weddell seals and ASM5032014v1 for leopard seals) of each species using bwa-mem v0.7.17-r1188 (Li, 2013), except for southern elephant seals, which were mapped to the northern elephant seal reference genome (mMirAng1.0.hap1) as described by Hoffman et al. (2024). This was justified by the higher quality and improved contiguity of the northern elephant seal assembly, as well as by the relatively recent divergence between the two species (0.4–4.0 million years ago, Arnason et al., 2006; Higdon et al., 2007). The resulting SAM files were converted to BAM format, sorted, filtered for duplicates and indexed with samtools 1.15.1 (Li et al., 2009). Variant calling was performed using bcftools mpileup v1.15.1 (Li, 2011) with a minimum mapping quality of 20 (min-MQ 20) and a minimum base quality of 20 (--min-BQ 20). The called variants were subsequently filtered using bcftools v1.15.1 (Li, 2011) and vcftools v0.1.16 (Danecek et al., 2011) to retain sites with a quality scores >100 and a mean (across individuals) sequencing depth of 5–40x. In addition, individual sites with sequencing depth below eight or above 80 were masked to exclude positions with unusually low or high coverage respectively. Finally, we applied a >33% missingness threshold across individuals.

### Sex scaffold removal

For each species, we identified and removed putative sex scaffolds prior to further analysis. For the southern elephant seal, sex scaffolds were removed as described by Hoffman et al. (2024). For the leopard seal, the four annotated sex scaffolds were directly excluded. For the crabeater and Weddell seals, where sex scaffold information was unavailable, putative sex-linked scaffolds were identified by extracting chain alignments between each target genome and the California sea lion (*Zalophus californianus*; GCA_900631625.1; Peart et al., 2021), a species with well characterised X and Y chromosomes. Alignments were obtained from a multi-species whole genome alignment of 13 pinniped species using Progressive Cactus v2.8.4 (Armstrong et al., 2020). Scaffolds with more than 20% of their sequence aligning to the California sea lion X or Y scaffolds were flagged as putative sex-linked regions. These candidates were then validated through visual inspection of ROH patterns, and the final set of sex-linked scaffolds was excluded from all downstream analyses.

### Genomic diversity, allele frequencies and inbreeding

We calculated per-site allele frequencies and nucleotide diversity (π) in 10 kb windows across all sites using vcftools v0.1.16 (Danecek et al., 2011). To quantify genomic inbreeding, we identified ROHs using bcftools v1.15.1 (Narasimhan et al., 2016), which applies a hidden Markov model to detect autozygous regions along the genome.Because ROH detection methods are commonly based on SNP data, their performance can be sensitive to the high density of variants in WGS datasets, which, despite quality filtering, is often accompanied by elevated genotyping error rates that can affect ROH identification. Specifically, false heterozygous calls can artificially break up ROHs, causing them to appear shorter or be missed entirely depending on the minimum length threshold. To mitigate this bias, we reduced SNP density prior to ROH calling by thinning the WGS datasets using VCFtools v0.1.16 (Danecek et al., 2011), retaining approximately one SNP every 3kb. This approach reduces the artificial fragmentation of ROHs while preserving a sufficient density of markers for robust ROH detection. After filtering, the number of SNPs retained was 771,127 for the crabeater seal, 697,462 for the leopard seal, 745,293 for the southern elephant seal and 690,569 for the Weddell seal. A minimum ROH length of 100kb was used to detect the true autozygous segments, and *F*_ROH_ was calculated for each individual as the proportion of the genome contained within ROHs.

### Demographic reconstruction

We inferred historical changes in *N*_e_ using SMC++ (v1.15.4; Terhorst et al., 2017). Autosomal VCF files were converted to SMC++ input format using smc++ vcf2smc, specifying the appropriate contig lengths. Analyses were performed assuming a mutation rate of μ = 7 × 10^-9^ mutations per site per generation (Peart et al., 2020) and a generation time (g) of 14.9 years (crabeater seal; Hückstädt, 2025), 10.4 years (leopard seal; Hückstädt, 2015a), 9.5 years (southern elephant seal; Hofmeyr, 2015) and 10.8 years (Weddell seal; Hückstädt, 2015b), taken from IUCN estimates. Uncertainty in *N*_*e*_ estimates was assessed using a block bootstrapping approach. Each chromosome was divided into 5 Mb segments, which were randomly resampled and concatenated to generate pseudochromosomes representing full-genome-size datasets. This was implemented using the publicly available script bootstrap_smcpp.py (available at https://github.com/popgenmethods/smcpp/issues/37). We generated 20 bootstrap replicates and SMC++ inference was repeated independently for each replicate to evaluate variability in demographic trajectories.

## Results and discussion

### Genome assembly

The final scaffolded and gap-closed genome assembly of the crabeater seal comprised 138 scaffolds with a total length of 2,444,645,479 bp (N50 = 99.1 Mb, N90 = 32.4 Mb; Figure 1). Gene completeness evaluated using BUSCO with the carnivora dataset identified 13,383 complete sequences, corresponding to 94.7% completeness, including 13,383 complete single-copy orthologs (92.3%), while 561 genes (3.9%) were missing. To further evaluate the completeness of the assembly, we aligned the Illumina and PacBio reads to the reference genome. The Illumina reads showed a mapping rate of 99.97% with a genome coverage of 99.98% and an average sequencing depth of 183x. The PacBio HiFi reads similarly achieved a mapping rate of 99.97%, with a genome coverage of 99.99% and an average sequencing depth of 44x.

**Figure 1.**
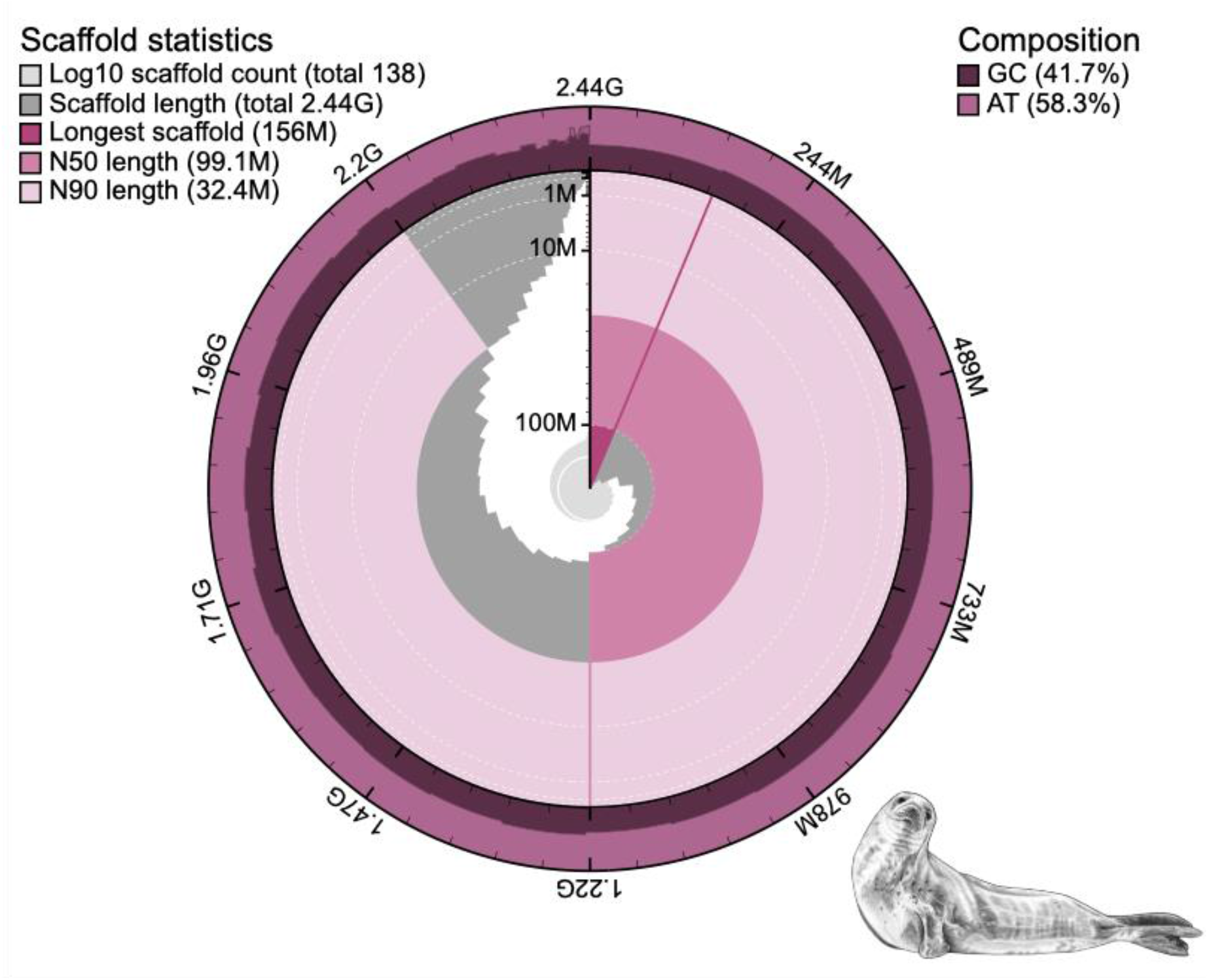
Snail plot summarizing the crabeater seal genome assembly. The innermost circle shows scaffold statistics scaled to the length of the longest scaffold, with the colours from dark to light purple indicating the longest scaffold, N50 and N90 lengths respectively, as shown in the legend. The light grey distribution shows the cumulative scaffold count on a log scale, while the dark grey distribution shows the scaffold length in descending order of size. GC and AT composition is shown in the outer circle. The seal illustration was created by Rebecca Carter (http://www.rebeccacarterart.co.uk) and is reproduced with her permission.

These assembly statistics compare favourably with other recent chromosomal pinniped reference genomes (see Supplementary table 1). Until recently, achieving this level of contiguity required chromatin conformation capture approaches such as HiC to link scaffolds, which are both costly and dependent on very high-quality input material. By contrast, the present assembly demonstrates that highly contiguous genomes can now be generated using a combination of Illumina and HiFi data without the need for HiC. Although not strictly chromosomal, the relatively small number of scaffolds (*n* = 138) represents a substantial improvement over earlier pinniped assemblies and showcases rapid advances in genome sequencing and assembly approaches.

### Genomic diversity and inbreeding

To evaluate the crabeater seal’s genomic diversity and place it into context, we computed the number of variant sites separately for each of the four species. Crabeater seals carried substantially more SNPs (61.5 million) than the other three species (∼12–16 million). In line with this, they also exhibited the highest π (0.0031, ± 0.00126 SD) compared to leopard (0.0010 ± 0.00054 SD), southern elephant (0.0016 ± 0.00101 SD) and Weddell seals (0.0009 ± 0.00064 SD; Figure 2A). Notably, these values are comparable to previous π estimates based on mitochondrial DNA or microsatellites (Bender et al., 2023; Curtis et al., 2009, 2011). Overall, these results suggest that crabeater seals have maintained high levels of standing genetic variation on a genome-wide scale, consistent with the expectation that large *N*_e_ promotes the accumulation and persistence of allelic diversity across generations.

**Figure 2.**
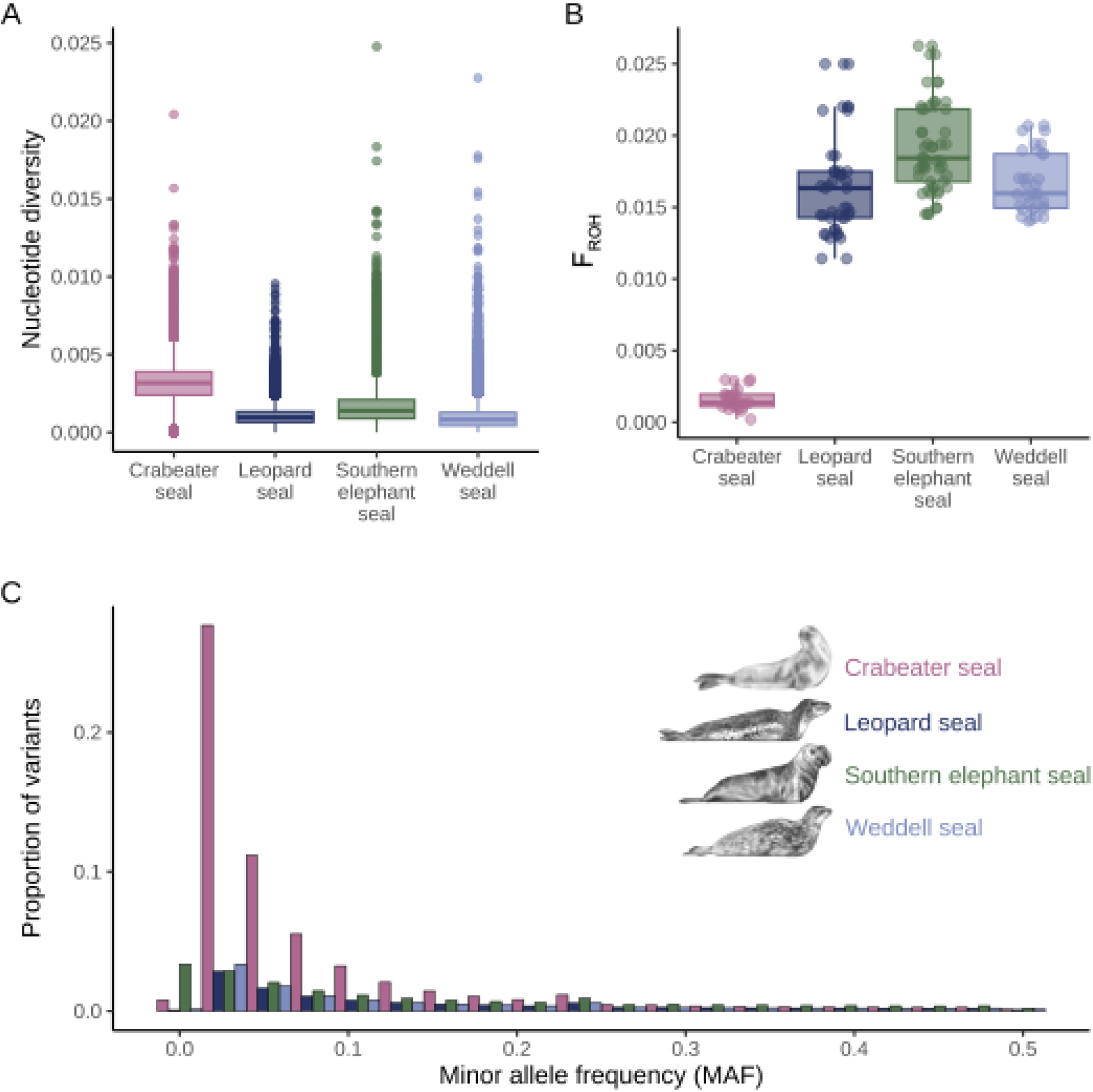
Nucleotide diversity, genome-wide inbreeding and allele frequency spectra in four Antarctic phocids: the crabeater seal, leopard seal, southern elephant seal and Weddell seal. (A) Nucleotide diversity (π) estimated in 10 kb non-overlapping windows across the genome; (B) individual inbreeding coefficients F_roh_ estimated from runs of homozygosity; (C) minor allele frequency (MAF) spectra showing the proportion of variants across frequency intervals. Raw data points are shown alongside boxplots; the thick centre lines represent median, the lower and upper hinges correspond to the first and third quartiles, respectively, and the whiskers represent largest and smallest value but no further than 1.5× the interquartile range. The bars in the barplots represent the proportion of variants falling within each frequency interval, with colours corresponding to species as indicated in the legend. The seal illustrations were created by Rebecca Carter (http://www.rebeccacarterart.co.uk) and are reproduced with her permission.

Patterns of genomic inbreeding differed markedly among species (Figure 2B). Crabeater seals exhibited consistently extremely low inbreeding (mean *F*_ROH_ = 0.002 ± 0.0007 SD). By contrast, the other three species showed *F*_ROH_ values that were one order of magnitude higher (Weddell seals: 0.017 ± 0.0021, leopard seals: 0.011 ± 0.0032, southern elephant seals: 0.019 ± 0.0032). Across all species, long ROHs (>5Mb), which are indicative of common ancestry within the last ∼10 generations, contributed relatively little to *F*_ROH_ (Supplementary figure 1). Instead, inbreeding was mostly attributable to short- and intermediate-length ROHs. These patterns suggest that genomic inbreeding in Antarctic phocids is shaped primarily by historical demography rather than recent consanguineous mating. Hence, the very low *F*_ROH_ of crabeater seals is consistent with a historically large *N*_e_.

### Allele frequency spectra and demographic histories

Because the SFS reflects past changes in population size, with an excess of rare alleles indicating population expansion and a flatter spectrum suggesting past bottlenecks, we inspected the SFS of each species. All four Antarctic phocids showed an excess of rare variants, largely reflected in negative Tajima’s D values and left-skewed allele frequency distributions (Figure 2C, Supplementary table 2). This pattern was most pronounced in the crabeater seal, where approximately half of all variants occurred at low frequency (MAF < 0.05: 48.96%). This enrichment of rare alleles in the crabeater seal is consistent with a non-equilibrium demographic history, most plausibly reflecting long-term population expansion, and it clearly distinguishes this species from the other three Antarctic phocids.

To investigate population size changes over time, we reconstructed historical *N*_e_ trajectories using SMC++, which implements a composite-likelihood framework based on the sequentially Markovian coalescent while leveraging information from multiple unphased genomes. The results are shown in Figure 3, while uncertainty in the estimates, assessed using a block bootstrapping approach, is shown in Supplementary figure 2. Crabeater seals exhibited consistently higher *N*_e_ over the past ∼1 million years and showed clear evidence of sustained population expansion (Figure 3), consistent with the excess of rare alleles observed in the SFS. Their *N*_e_ increased to a peak of around 1.6 million (95% bootstrap interval 0.59-0.66 million). By contrast, *N*_e_ was orders of magnitude lower for the other species, remaining below ∼50,000 for leopard seals, ∼60,000 for southern elephant seals and ∼95,000 for Weddell seals, and exhibiting smaller temporal fluctuations (Figure 3, Supplementary figure 2).

**Figure 3.**
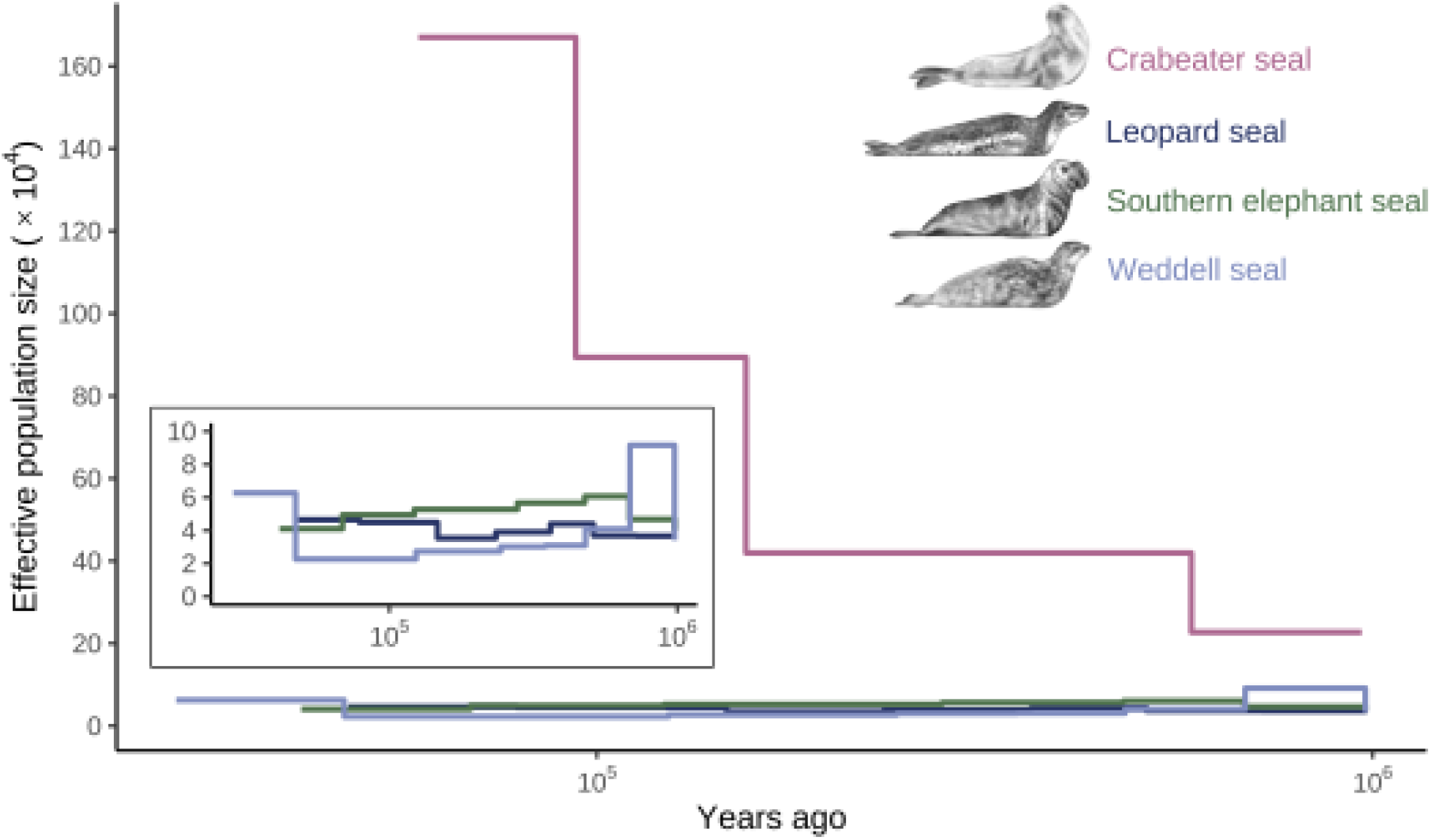
Historical effective population size (*N*_e_) trajectories inferred with SMC++. Coloured lines show the maximum-likelihood estimates of *N*_e_ based on whole-genome sequencing data from 20 individuals per species. Time is shown in years before present on a logarithmic scale. Analyses were restricted to autosomal chromosomes and assumed a mutation rate of μ = 0.7 × 10^−8^ mutations per site per generation (Peart et al., 2020) and species-specific generation times of 14.9 years (crabeater seal; Hückstädt, 2025), 10.4 years (leopard seal; Hückstädt, 2015a), 9.5 years (southern elephant seal; Hofmeyr, 2015) and 10.8 years (Weddell seal; Hückstädt, 2015b), taken from IUCN estimates. The inset shows the *N*_e_ estimates for leopard seal, southern elephant seal and Weddell seal on a rescaled y-axis (0-10 x 10^4^). The seal illustrations were created by Rebecca Carter (http://www.rebeccacarterart.co.uk/) and are reproduced with her permission.

The demographic expansion of crabeater seals is likely driven by their tight trophic coupling with Antarctic krill. Demographic reconstructions from krill genomes reveal a similar pattern of population expansion over the past ∼100,000 years, which is associated with cooler conditions during the late Pleistocene that increased the extent of sea ice habitat on which krill depend (Shao 2023). Crabeater seals also exhibit considerable behavioural plasticity in response to variation in krill availability, which may have helped to sustain their population growth by buffering against shorter-term fluctuations in prey availability (Hückstädt et al., 2012). This apparent demographic coupling between predator and prey aligns with patterns observed in another krill-dependent species, the Antarctic fur seal (*Arctocephalus gazella*) at South Georgia, where increased krill availability following the near extirpation of the great whales from the Southern Ocean was thought to have contributed to population recovery from commercial sealing (Hoffman et al. 2022), while more recent population declines since 2009 have been linked to reductions in krill (Forcada & Hoffman, 2014; Forcada et al., 2023). By contrast, the other three Antarctic phocids did not show such pronounced population expansions, likely because they exploit a broader range of prey species (Casaux et al., 2005; Bender et al., 2023; Woodman et al., 2024) whose populations may have been less sensitive to the specific environmental changes that favoured krill specialists.

## Conclusions

We assembled a high-quality crabeater seal genome and performed a comparative analysis of genomic diversity among Antarctic phocids. Consistent with population genetic theory, we observed high genome-wide diversity, low inbreeding and evidence for a large *N*_e_ and sustained population growth in crabeater seals. While most conservation genetic studies of wild populations have focused on small or declining populations, our results contribute to understanding the genomic consequences of very large population sizes. By investigating a thriving polar pinniped, this study contributes to a more complete understanding of the ecological and evolutionary processes that shape genetic diversity across the full spectrum of population sizes.

## Data availability

The raw reads of the genome assembly and the whole genome sequencing data from the four pinniped species is submitted to NCBI SRA BioProject No. PRJNA1478711 (https://www.ncbi.nlm.nih.gov/bioproject/PRJNA1478711/). The code associated with this study will be made publicly available upon acceptance.. Supplemental material available at G3 online.

## Acknowledgments

The authors thank the researchers, staff, and volunteers who contributed to the collection of the samples and field data used for this study. We are also grateful to Mallory Gulbranson for helping us with retrieving the crabeater seal samples from archived freezer collections and coordinating the shipment. We thank Ira Wachendorf and Ann-Christin Polikeit for carrying out DNA extractions and quality checks, as well as Rebecca Carter for providing the seal artwork.

## Author contributions

**crabeater seal:** TG carried out the fieldwork, collected and stored the 16 crabeater samples. The genome assembly sample and the remaining tissues used for whole genome resequencing of four crabeater seal samples were collected, stored, and provided by LAH, DPC and MT.

**Weddell seal:** JR, RG, and KM carried out the fieldwork, collected and stored the Weddell seal samples. EF and NC selected the samples for sequencing. EF and KM sourced the samples from archived freezer collections and coordinated their shipment.

**Southern elephant seal:** MEG, FG and SS contributed the Southern elephant seal samples used in this study.

**Leopard seal:** JF carried out the fieldwork and collected the leopard seal samples.

DLJV performed quality control on the samples prior to sequencing and shipment to the sequencing centre. TW was responsible for sequencing the samples at the West German Genome Center in Düsseldorf, except for the crabeater seals. JIH acquired funding and supervised the PhD student (BY). BY and JIH coordinated the study and drafted the manuscript. BY developed the methodology, performed the analyses and created the figures. All co-authors commented upon and approved the final manuscript.

## Supplementary material

**Table S1.** Assembly statistics of the crabeater seal reference genome compared to recent genome assemblies of the leopard seal, northern elephant seal, Antarctic fur seal and California sea lion.

|  | Crabeater seal | Leopard seal | Northern elephant seal | Antarctic fur seal | California sea lion |
| --- | --- | --- | --- | --- | --- |
| **Assembly ID** | LobCar_v1 | ASM5032014 v1 | mMirAng1.0. hap1 | arcGaz4_h1 | mZalCal1.pri. v2 |
| **Assembly level** | scaffold | scaffold | scaffold | scaffold | chromosome |
| **Genome size (bp)** | 2,444,645,479 | 2,454,374,544 | 2,430,305,188 | 2,527,999,884 | 2,409,685,272 |
| **Number of scaffolds** | 138 | 203 | 497 | 541 | 47 |
| **Scaffold N50 length** | 99,142,401 | 99,454,515 | 154,172,358 | 141,635,559 | 147,124,152 |
| **Scaffold N90 length** | 32,405,651 | 31,951,825 | 93,938,456 | 64,524,230 | 94,255,706 |
| **% BUSCO completeness** | 94.7<br>(carnivora_odb10) | 98.2<br>(carnivora_odb10) | 96.1<br>(mammalia_odb10) | 95.2<br>(carnivora_odb10) | 93.9<br>(mammalia_odb9) |

**Table S2.** Summary statistics of the allele frequency spectrum for four Antarctic phocids: the crabeater seal, leopard seal, southern elephant seal and Weddell seal. Proportions of singletons (minor allele count = 1) and doubletons (minor allele count = 2), and median Tajima’s D values (interquartile range) calculated in 10kb non-overlapping windows with at least five SNPs.

|  | **Crabeater seal** | **Leopard seal** | **Southern elephant seal** | **Weddell seal** |
| --- | --- | --- | --- | --- |
| **% singletons** | 44.1 | 20.8 | 10.3 | 24.3 |
| **% doubletons** | 17.3 | 11.5 | 7.3 | 11.5 |
| **Tajima's D** | -1.76 (-1.92, -1.57) | -0.38 (-0.86, 0.1) | 0.03 (-0.51, 0.56) | -0.51 (-0.1,-0.02) |

**Figure S1.**
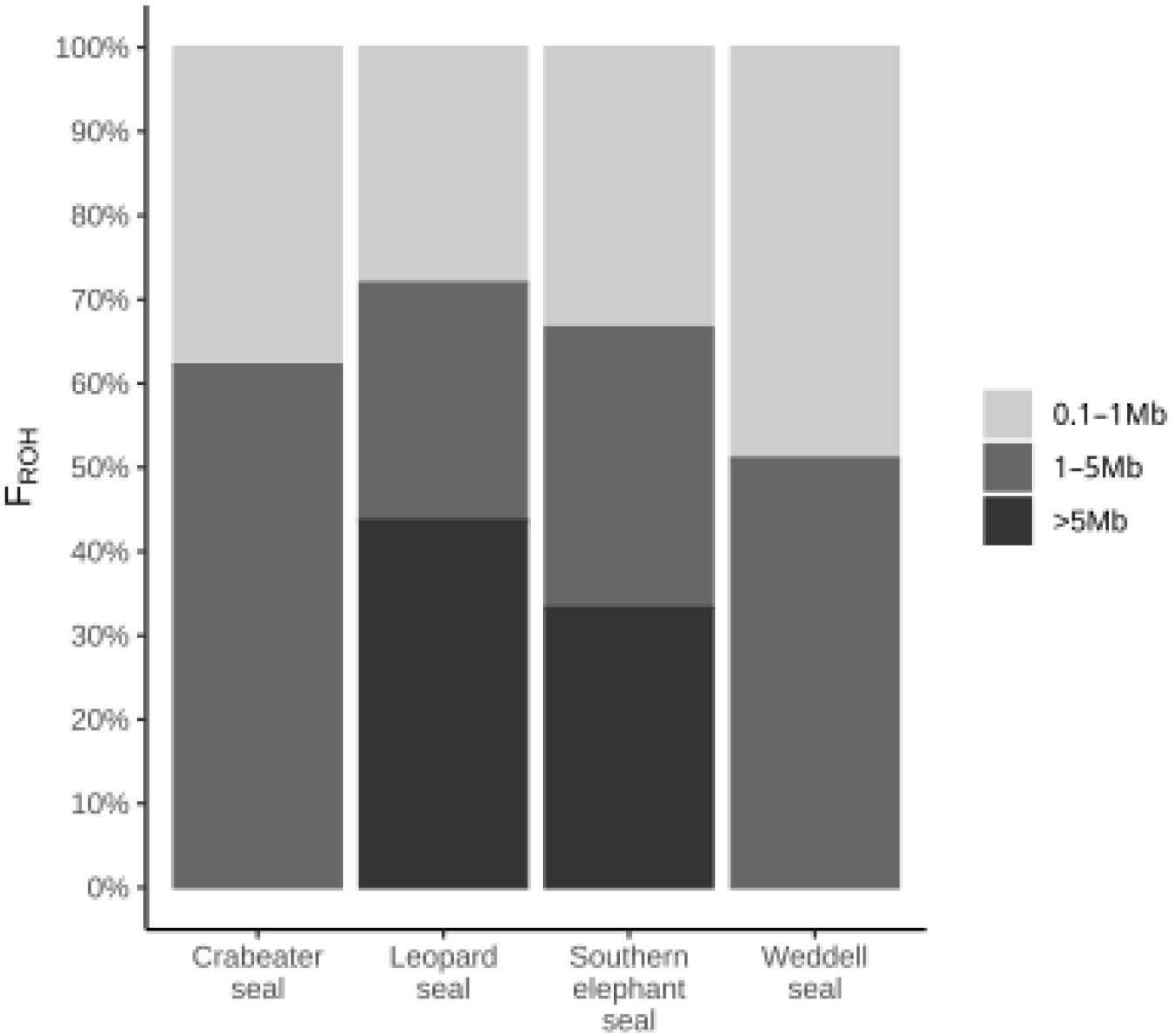
Genome-wide inbreeding composition across ROH length categories for four Antarctic phocids: the crabeater seal, leopard seal, southern elephant seal and Weddell seal. Individual inbreeding coefficients (*F*_ROH_) estimated from runs of homozygosity (ROHs) are grouped into three length categories: 0.1–1 Mb, 1–5 Mb and >5 Mb. The stacked bars show the mean proportion of the total *F*_*ROH*_ attributable to each ROH length class per species. Short ROHs reflect more ancient shared ancestry, whereas long ROHs indicate more recent shared ancestry.

**Figure S2.**
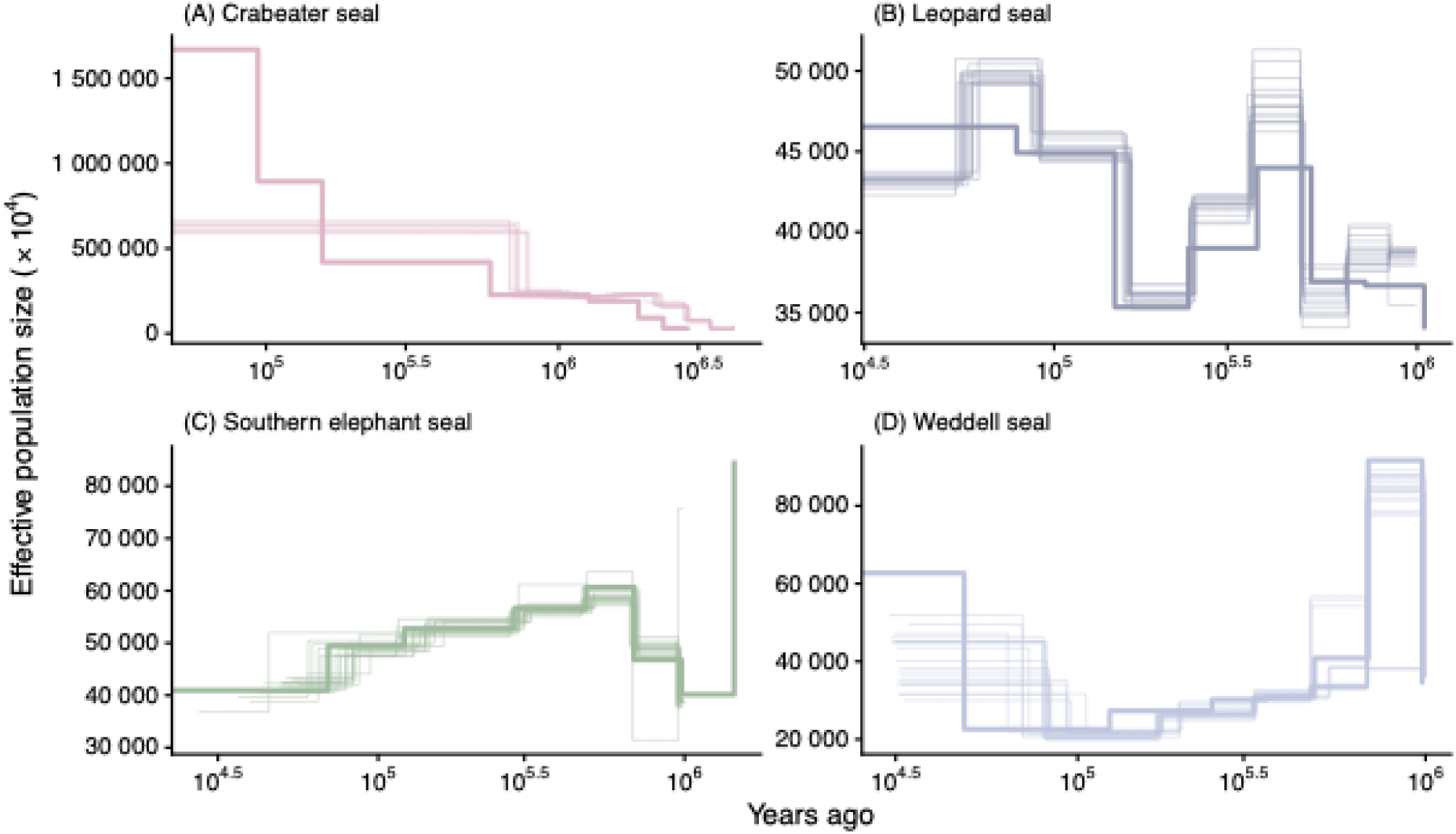
Uncertainty in historical effective population size (*N*_e_) trajectories inferred using SMC++. Bold coloured lines show the maximum-likelihood estimates of *N*_e_ based on whole-genome sequencing data from 20 individuals per species, while the thin lines show *N*_e_ trajectories from 20 block bootstrap replicates generated by resampling 5 Mb genomic segments. Time is shown in years before present on a logarithmic scale. Analyses were restricted to autosomal chromosomes and assumed a mutation rate of μ = 0.7 × 10^−8^ mutations per site per generation (Peart et al., 2020) and species-specific generation times of 14.9 years (crabeater seal; Hückstädt, 2025), 10.4 years (leopard seal; Hückstädt, 2015a), 9.5 years (southern elephant seal; Hofmeyr, 2015) and 10.8 years (Weddell seal; Hückstädt, 2015b), taken from IUCN estimates.

